# DNA barcoding affirms the presence of invasive *Parachanna obscura* and new records of some species in the Mweru-Luapula fishery

**DOI:** 10.1101/2024.02.11.579834

**Authors:** Bornwell Seemani, Cyprian Katongo, Paulette Bloomer, Arrie W. Klopper, Carel J. Oosthuizen

**Author notes:** Corresponding author (BS).

## Abstract

DNA barcoding has recently been instrumental in identifying both invasive and undetected species in aquatic environments. This study aimed at analysing collected fish fin clips to ascertain the identity of native species and the *Parachanna* species that have invaded the Mweru-Luapula (ML) fishery of Zambia. The identification process was carried out through field phenotypic analysis using species guides and DNA barcoding, with the mitochondrial DNA (mtDNA) cytochrome C oxidase 1 (COI) gene fragment. Of the 28 specimens for which DNA was successfully PCR amplified, five matched the reference sequences of species and 22 matched the reference sequences of genera on the NCBI GenBank. Five unexpected species, namely *Oreochromis niloticus, Coptodon zillii, Mormyrus kannume, Thoracochromis buysi* and *Tylochromis polylepis* were identified. The study further affirmed the presence of invasive *Parachanna obscura* in the fishery and its interconnected water bodies. *Parachanna obscura* invaded the fishery through annual flooding from aquacultural facilities in the Democratic Republic of Congo (DRC). There is a need to investigate this invasion further by using large sample sizes and also by applying the gonadosomatic index (GSI) to determine invasive species occupancy and impact on native species throughout the fishery. This study provides a platform for further detailed taxonomic verification and species inventory of the entire ML fishery. This will facilitate the development of a viable and sustainable strategy to appropriately curb the impact of invasive species, and will thus contribute to the conservation of the ML aquatic biodiversity.

## Introduction

The application of DNA barcoding in studying freshwater fishes and their vast ecosystems avails conservationists a wide set of information, which includes, inter alia rapid identification of invasives, aid to understanding of threatened species and biology of cryptic species [1–3]. DNA barcoding is a fast-growing tool that has been incorporated in the management and monitoring of various ecosystems across the globe [1,3,4). The technique is robust, reliable and the most accurate method of fish identification [5] down to the species level [6], and can also be applied in population delineation studies [7]. DNA barcodes provide a reference database to detect and identify invasive species within impacted communities [8]. This tool has gained global popularity over conventional and traditional methods because of its speed and efficiency in identifying fish species [4].

For the past few years, tremendous progress has been made in the application of DNA barcodes in the development of both aquaculture and capture fisheries. However, policies and the practical utilisation of DNA barcodes has remained stagnant in most third world countries [3]. DNA barcodes cannot replace taxonomists and conventional species monitoring methods, but instead, they can complement the already existing field monitoring tools [1]. Management of fisheries resources is dependent upon valid and effective species identification [4]. This efficiency in identification consequently contributes to biodiversity conservation [9] and is an essential component in all science disciplines [10]. Species diagnoses drive biological research and development to prevent taxonomic proficiency from collapsing [2]. DNA barcoding has contributed immensely to the discovery of unidentified species, even in well-studied ecosystems [1]. Thus, genetic identification plays a vital part in the sustainable management of aquatic species under small-scale fisheries [5].

Fish fin clips obtained from either dead or live fish are being extensively used as a source for extracting genomic DNA [11]. Fin clipping is among the old techniques that has been applied for marking fish [12] as this technique exerts minimal deleterious effects on live fish [13]. It also requires nominal equipment to execute [14], accessed easily and delivers good quality DNA for molecular analyses [15]. By using fish fin clips, Holmes et al. [16] were able to identify specimens to species level with short sequences of only 398 bp of DNA barcoding fragments. Mehrdad Hajibabaei et al. (2006) [17]16 also effectively identified a specimen based on 109 bp of the cytochrome C oxidase 1 (COI) barcoding region.

The mitochondrial DNA (mtDNA) cytochrome C oxidase 1 (COI) gene fragment is utilised as a species identifying marker [18,19]. Hebert et al. [2] recommended COI to be ideal for animals, including fish [15,20]. It has been found that COI provides a dependable and cost-effective approach to species identification [2]. The COI is regularly used as a universal barcode due to the level of sequence conservation linked to the function of its protein product [21]. Arabi et al. [22] have also stressed the appropriateness of the conserved COI in assessing phylogenetic relationships. The gene can be used to identify species from sequences of about 600 bp long [2,19,23]. DNA barcoding using COI has become an extensive and dependable marker with the rapidly growing Barcode of Life Data System (BOLD) [24]. The COI DNA fragment has also been applied in the study of invasive species, including the freshwater *Parachanna obscura* as demonstrated by Serrao et al. [8] and Conte-Grand et al. [25].

*Parachanna obscura* is an African snakehead fish belonging to the Channidae family. Snakeheads are species of vast economic importance to both fisheries and aquaculture due to their high nutritional value and high market demand [18]. They are preferred for consumption in many African countries, hence their huge potential as part of aquaculture development [26]. On the other hand, *Oreochromis* species are part of the over 2,500 cichlids spread across the three continents of Africa, America and Asia [27]. A large component of cichlid diversity has evolved within the last 10 million years [28]. Cichlids are also renowned for adaptive radiations in East Africa [29]. *Oreochromis macrochir* (Boulenger, 1912), the greenhead tilapia, is a common detritivorous cichlid [30], that is increasingly becoming a commercial and recreational candidate for aquaculture development in Zambia [31]*. Oreochromis* species are native to the Mweru-Luapula (ML) fishery and other major fisheries of the country.

The ML fishery, located in the northern region of Zambia is diverse, consisting of approximately 135 freshwater species from over 16 families [32]. Fish stock assessment of the fishery depends on fish data derived from traditional methods, which include gill net surveys (GNSs), catch assessment surveys (CASs), frame surveys (FSs), market statistics and limnological studies [33,34]. These independent fishery surveys are laborious, costly and cannot readily capture data on cryptic species. To-date, no comprehensive DNA barcoding studies have been done on freshwater fishes within the ML fishery. In this study, we made use of DNA barcodes to assess whether the species that has invaded the ML fishery and its interconnected water bodies is *P. obscura* [8] or a potentially new *Parachanna* species (*Pa*. sp. D. R. Congo, BIN AAF7843) observed by Conte-Grand et al. [25]. *Parachanna* invaded the fishery through annual flooding from DRC [35,36]. This study compared traditional taxonomic identification to the use of DNA barcodes and analysed species relationships using phylogenetic trees.

## Materials and methods

### Study area

The study was conducted in the ML fishery of Zambia, located at 8° 28’ – 9° 31’ S, and 28° 20’ – 29° 20’ E [37]. Samples were collected from 11 randomly selected locations (of 18 possible sampling localities) within the ML fishery. Samples were obtained after fish catches from gill net surveys (GNS) and local fishers [26] were recorded. These sampling localities were distributed throughout the four strata of the fishery (Fig 1). The 11 sampling sites were Puta-Abinala (Stratum 1), Kenani and Mwatishi (Stratum II), Kilwa Islands, Chisenga Islands-Muku lagoon, Mofwe Lagoon (Stratum III) and Chipita-Filumba Lagoon, Luapula River (LR I), Luche Lagoon-Lukwesa, Luapula River-Lukwesa (LR II) and Nshinda stream (Stratum 4).

**Fig 1.**
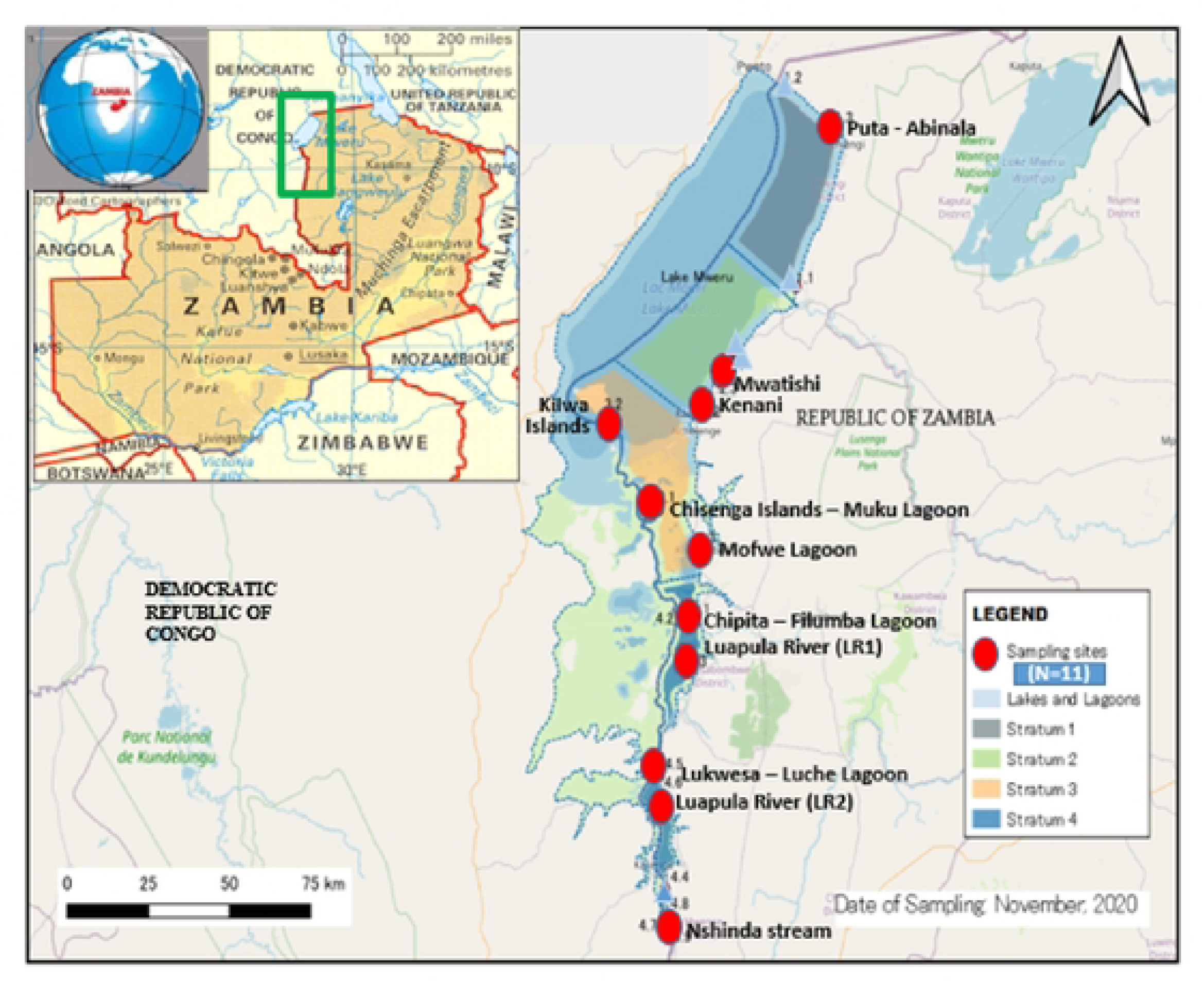
The Mweru-Luapula fishery showing the location of 11 sampling sites where fin clips (designated by red dots) were collected. Inset is the Map of Zambia, depicting the location (green rectangle) of the fishery.

### Sample collection

Sampling was conducted using two engine-driven fibre research boats that carried the research team and two fleets of bottom and top set nets. The fleets were set overnight in a single haul for a soak time ranging from 8-14 hours in all targeted sampling sites. The bottom set fleet was 650 m long, consisting of 13 nets (25, 37, 50, 63, 76, 89, 102, 114, 127, 140, 152, 165 and 178 mm). The top set fleet was 600 m long, consisting of 12 nets (25, 37, 50, 63, 76, 89, 102, 114, 127, 140, 152 and 165 mm). Average depths for bottom set and top set fleets were 2.5 m and 2.7 m respectively (S1 Table). Both fleets were mounted at a set hanging ratio of 0.5, following the DoF – Central Fisheries Research Institute (CFRI) guidelines and protocol. All the nets were multifilament, except for one 76.2 mm monofilament nylon twine on the top set fleet. The two fleets were fished as separate gears and set perpendicular to the shoreline. Garmin Etrex 10 – Geographical positioning system (GPS) was used to record the precise sampling locations. Sampled fish specimens were morphologically identified to species level using identity guides by Skelton [38] and Utsugi and Mazingaliwa [39] (Table 1).

**TABLE 1.**
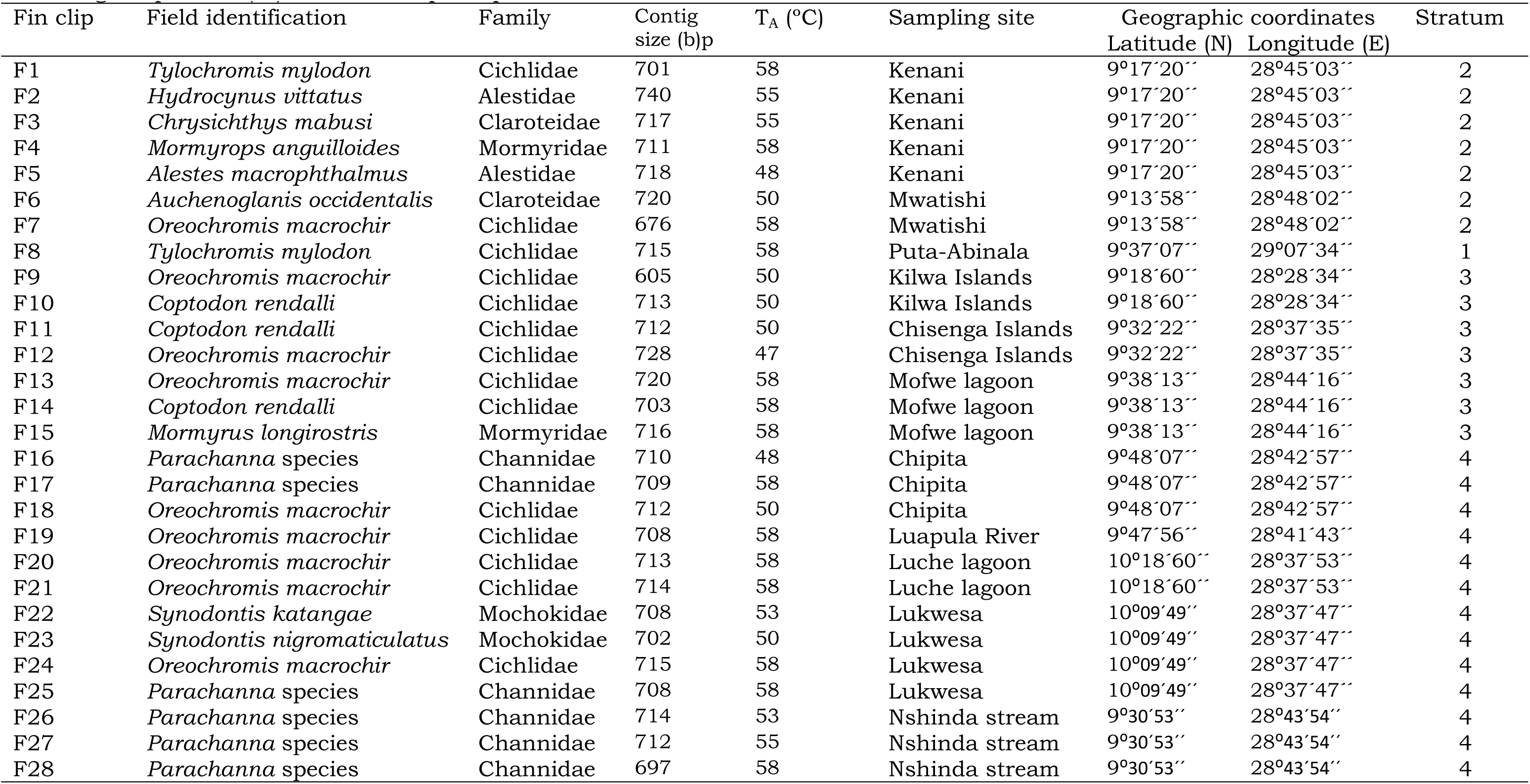
Details of the fin clips collected (F1 – F28), corresponding field identification (species and family), sampling site, sampling site coordinates and the stratum represented within the Mweru-Luapula fishery are provided. The associated contig size and primer annealing temperature(*T*_A_) for each sample is provided.

| Fin clip | Field identification | Family | Contig size (b)p | $T_A$ (°C) | Sampling site | Geographic coordinates | | Stratum |
| --- | --- | --- | --- | --- | --- | --- | --- | --- |
|  |  |  |  |  |  | Latitude (N) | Longitude (E) |  |
| F1 | *Tylochromis mylodon* | Cichlidae | 701 | 58 | Kenani | 9°17'20" | 28°45'03" | 2 |
| F2 | *Hydrocynus vittatus* | Alestidae | 740 | 55 | Kenani | 9°17'20" | 28°45'03" | 2 |
| F3 | *Chrysichthys mabusi* | Claroteidae | 717 | 55 | Kenani | 9°17'20" | 28°45'03" | 2 |
| F4 | *Mormyrops anguilloides* | Mormyridae | 711 | 58 | Kenani | 9°17'20" | 28°45'03" | 2 |
| F5 | *Alestes macrophthalmus* | Alestidae | 718 | 48 | Kenani | 9°17'20" | 28°45'03" | 2 |
| F6 | *Auchenoglanis occidentalis* | Claroteidae | 720 | 50 | Mwatishi | 9°13'58" | 28°48'02" | 2 |
| F7 | *Oreochromis macrochir* | Cichlidae | 676 | 58 | Mwatishi | 9°13'58" | 28°48'02" | 2 |
| F8 | *Tylochromis mylodon* | Cichlidae | 715 | 58 | Putu-Abinala | 9°37'07" | 29°07'34" | 1 |
| F9 | *Oreochromis macrochir* | Cichlidae | 605 | 50 | Kilwa Islands | 9°18'60" | 28°28'34" | 3 |
| F10 | *Coptodon rendalli* | Cichlidae | 713 | 50 | Kilwa Islands | 9°18'60" | 28°28'34" | 3 |
| F11 | *Coptodon rendalli* | Cichlidae | 712 | 50 | Chisenga Islands | 9°32'22" | 28°37'35" | 3 |
| F12 | *Oreochromis macrochir* | Cichlidae | 728 | 47 | Chisenga Islands | 9°32'22" | 28°37'35" | 3 |
| F13 | *Oreochromis macrochir* | Cichlidae | 720 | 58 | Mofwe lagoon | 9°38'13" | 28°44'16" | 3 |
| F14 | *Coptodon rendalli* | Cichlidae | 703 | 58 | Mofwe lagoon | 9°38'13" | 28°44'16" | 3 |
| F15 | *Mormyrus longirostris* | Mormyridae | 716 | 58 | Mofwe lagoon | 9°38'13" | 28°44'16" | 3 |
| F16 | *Parachanna species* | Channidae | 710 | 48 | Chipita | 9°48'07" | 28°42'57" | 4 |
| F17 | *Parachanna species* | Channidae | 709 | 58 | Chipita | 9°48'07" | 28°42'57" | 4 |
| F18 | *Oreochromis macrochir* | Cichlidae | 712 | 50 | Chipita | 9°48'07" | 28°42'57" | 4 |
| F19 | *Oreochromis macrochir* | Cichlidae | 708 | 58 | Luapula River | 9°47'56" | 28°41'43" | 4 |
| F20 | *Oreochromis macrochir* | Cichlidae | 713 | 58 | Luche lagoon | 10°18'60" | 28°37'53" | 4 |
| F21 | *Oreochromis macrochir* | Cichlidae | 714 | 58 | Luche lagoon | 10°18'60" | 28°37'53" | 4 |
| F22 | *Synodontis katangae* | Mochokidae | 708 | 53 | Lukwesa | 10°09'49" | 28°37'47" | 4 |
| F23 | *Synodontis nigromaticulatus* | Mochokidae | 702 | 50 | Lukwesa | 10°09'49" | 28°37'47" | 4 |
| F24 | *Oreochromis macrochir* | Cichlidae | 715 | 58 | Lukwesa | 10°09'49" | 28°37'47" | 4 |
| F25 | *Parachanna species* | Channidae | 708 | 58 | Lukwesa | 10°09'49" | 28°37'47" | 4 |
| F26 | *Parachanna species* | Channidae | 714 | 53 | Nshinda stream | 9°30'53" | 28°43'54" | 4 |
| F27 | *Parachanna species* | Channidae | 712 | 55 | Nshinda stream | 9°30'53" | 28°43'54" | 4 |
| F28 | *Parachanna species* | Channidae | 697 | 58 | Nshinda stream | 9°30'53" | 28°43'54" | 4 |

Twenty-eight pectoral fin clips were collected, representing individual fish species from six families, including Cichlidae (n=14), Channidae (n=6), Alestidae (n=2), Claroteidae (n=2), Mochokidae (n=2) and Mormyridae (n=2) (Table 1). Fin clip collection was performed during November 2020. Clipping of pectoral fin [16,40] from dead fish was done with a sterilized pair of scissors and thereafter stored in sterilized 1.8 ml serum vials filled with 95% ethanol and kept at −20°C until the laboratory analysis. A pair of scissors/scalpel and forceps were subjected to 30% bleach and rinsed in distilled water between individual fish and sampling sites. Each fin clip served as a subsample of collected specimens. Due to limited reagents, only a few representative specimens were wholly preserved in formalin and are being kept at the University of Zambia.

### DNA extraction and amplification by PCR

Total genomic DNA was extracted from 10% suspension of fin clips (<1 mm^2^) using the DNeasy Blood and Tissue Kit (Qiagen) according to the manufacturer’s protocols. DNA quantity and quality was assessed using a Nanodrop 2000 spectrophotometer (Thermo Fisher Scientific Inc). The COI fragments for each sample was PCR amplified (Thermo Fisher Scientific Inc) in 25 μl total reaction volumes using SuperTherm *Taq* according to the manufacturer’s procedure. The reaction contained 50 ng template DNA, 2.5 mM MgCl_2_, 1 x buffer, 0.2 mM dNTPs (for all four nucleotides) and 1 U of Super-Therm *Taq* polymerase (Separation Scientific), 0.01 mM of each fish primer, VF2-t1 (5’ TGT AAA ACG ACG GCC AGT CAA CCA ACC ACA AAG ACA TTG GCA C 3’) and FishR2-t1 (5’ CAG GAA ACA GCT ATG ACA CTT CAG GGT GAC CGA AGA ATC AGA A ‘3) [41] and ddH_2_O to make up the final volume to 25 μl. The PCR programme followed consisted of initial denaturation for 4 min at 94°C, 35 cycles of 30s at 94°C, primer annealing for 30s at 47°C – 58°C (Table 1) and 30s of elongation at 72°C, and a final elongation step of 7 min at 72°C [42,43]. PCR products were tested for amplification success via gel electrophoresis through a 1% (w/v) agarose gel stained using GelRed (Anatech Instruments PTY LTD). Negative controls were performed in the extraction and amplification steps to assess any potential contamination through a blank containing only buffer. Each gel contained a 100 bp size standard (Inqaba Biotechnical Industries PTY LTD) [44] to assess amplicon size and concentration.

### Sequencing and alignment

Successful PCR amplicons were precipitated for sequencing by adding 30 μl of 100% ethanol, 5.5 μl of ddH_2_O and 1 μl of 3 M sodium acetate to the sample. The mixture was centrifuged at 13,000 rpm for 15 min, before adding 90 μl of freshly made up 70% ethanol. The mixture was centrifuged again for 10 min at 13,000 rpm, before air drying the DNA on a heat block at 45°C after removing the supernatant. The dried pellets were then resuspended in 10 μl of dH_2_O before the purified PCR products were sequenced in both forward and reverse directions. Sequencing reactions were performed using 0.125 μl of ABI PRISM BigDye^®^ Terminator V3.1 Cycle Sequencing Kit (Applied Biosystems), 5x sequencing Buffer, 3.2 pmol of FishR2-t1 (reverse) or VF2-t1 (forward) primer, 50 ng of DNA and ddH_2_O to make up the final volume to 10 μl. The cycle sequencing programme followed consisted of initial denaturation for 10s at 96°C, 25 cycles of 5s at 94°C, primer annealing for 30s at 55-58°C and 4 min of elongation at 60°C, and kept at 4°C. Thereafter, sequencing reactions were performed using the ABI 3500xl (Applied Biosystems).

The raw sequencing chromatopherograms for each of the 28 mtDNA COI sequences were visually inspected and manually edited where needed. Forward and reverse sequences for each sample was aligned using the CLC Main Workbench V 21.0.5 (QIAGEN). The consensus sequences (contigs) for each sample were aligned to each other using ClustalX V 2.1 [45] with all mutational difference observed again confirmed by referring to the original chromatopherogram. These contigs were then individually used in a Basic Local Alignment Search Tool (BLAST) [46] search on the National Centre for Biotechnology Information (NCBI) platform to find the best matches for all 28 sequences on the GenBank database [26].

### Phylogenetic analysis and sequence divergence

To assess the phylogenetic relationships for taxa sequenced in this study, 57 additional COI sequences were retrieved from GenBank. This included five outgroup taxa, GenBank sequences of *C. citharinus* HQ927822.1 [47], *Pristolepis fasciata* [48], *Acanthocleithron chapini* KT192951.1 [49], *Hepsetus odoe* HM882992.1 [50], *Bagrus bajad* MK335908.1 [51] and *Marcusenius senegalensis* HM882735.1 [52] were employed as outgroups for cichlids, channids, mochokids, alestids, claroteids and mormyrids for the analyses, respectively. The rest of the sequences represented six families: Cichlidae, Channidae, Mochokidae, Alestidae, Claroteidae and Mormyridae (S2 Table). Both the retrieved and sequences generated from this study (contigs) were aligned and converted in MEGA 11 [53] for the construction of phylogenetic trees using Maximum Likelihood (ML) [54,55], with 1000 bootstrap repeats. As is standard in DNA barcoding studies, the Kimura-2-Parameter model was used to calculate the sequence divergence among the sequences and species [18,56].

## Results

### Confirmation of species identity

A total of 28 cytochrome C oxidase I (COI) sequences (all >650 bp fragments) were obtained, belonging to 12 morphologically identified fishes (Table 2). All these sequences were individually used in a BLAST search on NCBI.

**TABLE 2.** BLAST results of 28 samples obtained from the Mweru-Luapula fishery, sequenced for the COI barcoding gene for fish species identification. F1 – F28 are the samples collected with their associated field identification, BLAST top hit, family, total score, percent identity (*I* %), query cover, accession length and number.

| Fin clip | Field identification | BLAST Top Hit | Family | Total score | *I* % | Query cover % | Accession length bp | Accession number |
| --- | --- | --- | --- | --- | --- | --- | --- | --- |
| F1 | *Tylochromis mylodon*<sup>a</sup> | *Tylochromis polylepis*<sup>a</sup> | Cichlidae | 1254 | 99.00 | 99 | 16976 | AP009509.1 |
| F2 | *Hydrocynus vittatus* | *Hydrocynus vittatus* | Alestidae | 1005 | 96.69 | 81 | 609 | KX186117.1 |
| F3 | *Chrysichthys mabusi* | *Chrysichthys mabusi* | Claroteidae | 1086 | 99.83 | 82 | 591 | HG803412.1 |
| F4 | *Mormyrops anguilloides* | *Mormyrops anguilloides* | Mormyridae | 1186 | 99.54 | 91 | 651 | KT193238.1 |
| F5 | *Alestes macrophthalmus*<sup>b</sup> | *Thoracochromis buysi*<sup>b</sup> | Cichlidae | 1177 | 99.23 | 90 | 652 | JN312262.1 |
| F6 | *Auchenoglanis occidentalis* | *Auchenoglanis occidentalis* | Bagridae | 1194 | 99.69 | 90 | 652 | HM880259.1 |
| F7 | *Oreochromis macrochir*<sup>c</sup> | *Oreochromis niloticus*<sup>c</sup> | Cichlidae | 1194 | 100.00 | 95 | 655 | KT307766.1 |
| F8 | *Tylochromis mylodon*<sup>a</sup> | *Tylochromis polylepis*<sup>a</sup> | Cichlidae | 1256 | 99.00 | 98 | 16976 | AP009509.1 |
| F9 | *Oreochromis macrochir*<sup>c</sup> | *Oreochromis niloticus*<sup>c</sup> | Cichlidae | 1064 | 99.83 | 95 | 655 | KT307766.1 |
| F10 | *Coptodon rendalli*<sup>d</sup> | *Coptodon zillii*<sup>d</sup> | Cichlidae | 1219 | 100.00 | 92 | 660 | KJ938170.1 |
| F11 | *Coptodon rendalli*<sup>d</sup> | *Coptodon zillii*<sup>d</sup> | Cichlidae | 1219 | 100.00 | 92 | 660 | KJ938170.1 |
| F12 | *Oreochromis macrochir*<sup>c</sup> | *Oreochromis niloticus*<sup>c</sup> | Cichlidae | 1205 | 99.85 | 89 | 655 | KT307766.1 |
| F13 | *Oreochromis macrochir*<sup>c</sup> | *Oreochromis niloticus*<sup>c</sup> | Cichlidae | 1210 | 100.00 | 90 | 655 | KT307766.1 |
| F14 | *Coptodon rendalli*<sup>d</sup> | *Coptodon zillii*<sup>d</sup> | Cichlidae | 1214 | 99.85 | 93 | 660 | KJ938170.1 |
| F15 | *Mormyrus longirostris*<sup>e</sup> | *Mormyrus kannume*<sup>e</sup> | Mormyridae | 1147 | 98.46 | 90 | 651 | MT523649.1 |
| F16 | *Parachanna species* | *Parachanna obscura* | Channidae | 1199 | 99.85 | 91 | 652 | MK074551.1 |
| F17 | *Parachanna species* | *Parachanna obscura* | Channidae | 1199 | 99.85 | 91 | 652 | MK074551.1 |
| F18 | *Oreochromis macrochir*<sup>c</sup> | *Oreochromis niloticus*<sup>c</sup> | Cichlidae | 1205 | 99.85 | 91 | 655 | KT307766.1 |
| F19 | *Oreochromis macrochir*<sup>c</sup> | *Oreochromis niloticus*<sup>c</sup> | Cichlidae | 1205 | 99.85 | 92 | 655 | KT307766.1 |
| F20 | *Oreochromis macrochir*<sup>c</sup> | *Oreochromis niloticus*<sup>c</sup> | Cichlidae | 1205 | 99.85 | 91 | 655 | KT307766.1 |
| F21 | *Oreochromis macrochir*<sup>c</sup> | *Oreochromis niloticus*<sup>c</sup> | Cichlidae | 1205 | 99.85 | 91 | 655 | KT307766.1 |
| F22 | *Synodontis katangae*<sup>f</sup> | *Synodontis nigromaculata*<sup>f</sup> | Mochokidae | 1245 | 99.85 | 95 | 677 | HF565919.1 |
| F23 | *Synodontis nigromaculata* | *Synodontis nigromaculata* | Mochokidae | 1240 | 99.70 | 96 | 677 | HF565919.1 |
| F24 | *Oreochromis macrochir*<sup>c</sup> | *Oreochromis niloticus*<sup>c</sup> | Cichlidae | 1205 | 99.85 | 91 | 655 | KT307766.1 |
| F25 | *Parachanna species* | *Parachanna obscura* | Channidae | 1199 | 99.85 | 92 | 652 | MK074551.1 |
| F26 | *Parachanna species* | *Parachanna obscura* | Channidae | 1199 | 99.85 | 91 | 652 | MK074551.1 |
| F27 | *Parachanna species* | *Parachanna obscura* | Channidae | 1199 | 99.85 | 91 | 652 | MK074551.1 |
| F28 | *Parachanna species* | *Parachanna obscura* | Channidae | 1199 | 99.85 | 93 | 652 | MK074551.1 |

**Mismatches between field identification and the GenBank database**
<sup>a</sup>*Tylochromis mylodon* as *Tylochromis polylepis*; <sup>b</sup>*Alestes macrophthalmus* as *Thoracochromis*; <sup>c</sup>*Oreochromis macrochir* as *Oreochromis niloticus*;
<sup>d</sup>*Coptodon rendalli* as *Coptodon zillii*; <sup>e</sup>*Mormyrus longirostris* as *Mormyrus kannume* and <sup>f</sup>*Synodontis katangae* as *Synodontis nigromaculata*

The “Top Hit” percentage identification match was taken to be the ‘best match’; in most instances the estimate of identity had scores above 99%, except for two samples, F2 (*Hydrocynus vittatus*) at 96.69% and F15 (*Mormyrus kannume*) at 98.46% (Table 2). The E-value for all samples was zero. A single specimen was recorded as a complete mismatch (*Thoracochromis buysi* identified as *Alestes macrophthalmus* based on field phenotypic observations), with 16 specimens matching the correct genera, but wrong species. Meanwhile 11 specimens showed a 100% match to a reference sequence (Table 2).

The BLAST search revealed the presence of invasive *P. obscura* (in stratum IV) and *Oreochromis niloticus* (in strata III and IV). All species classified as *O. macrochir* during field identification matched *O. niloticus* using the BLAST search on GenBank (F7 & F13 –100.00%; F12, F18, F19, F20, F21 & F24 – 99.85%; and F9 – 99.83%).

### Cichlidae

All nine *Oreochromis* specimens (F7, F9, F12, F13, F18, F19, F20, F21 & F24) formed a well-defined group, with a ‘Top Hit’ or best match to *O. niloticus* KT307766.1 with similarity identities of 100% (F7 & F13), 99.85% (F12, F18, F19, F20, F21 & F24) and 99.83% (F9) (Table 2). The *Oreochromis* tree showed two separate lineages for the oreochromine cichlids, with the second one comprising *Oreochromis* species 1 and *Oreochromis* species 3 not native to the ML fishery. Specimens F1 and F8 with similarity identity of 99.00% clustered with *T. polylepis* AP009509.1, whereas F5 with similarity identity of 99.23% clustered with *T. buysi* JN312262.1. From the phylogenetic analyses it is clear that the *Coptodon* species consist of at least two distinct lineages. The three *Coptodon* specimens (F10, F11 & F14) were assigned to a single species clade. Specimens F10 and F11 had a 100% similarity to *C. zilli* KJ938170.1 using the maximum likelihood model in MEGA 11. The analysis further classified all three specimens (F10, F11 & F14) as closely related to both *C. zilli* KJ938170.1 and *C. rendalli* MG438461.1. The sequence divergence of 0.3% shows specimens representing *C. zilli* and *C. rendalli* could belong to the same species (Fig 2).

**Fig 2.**
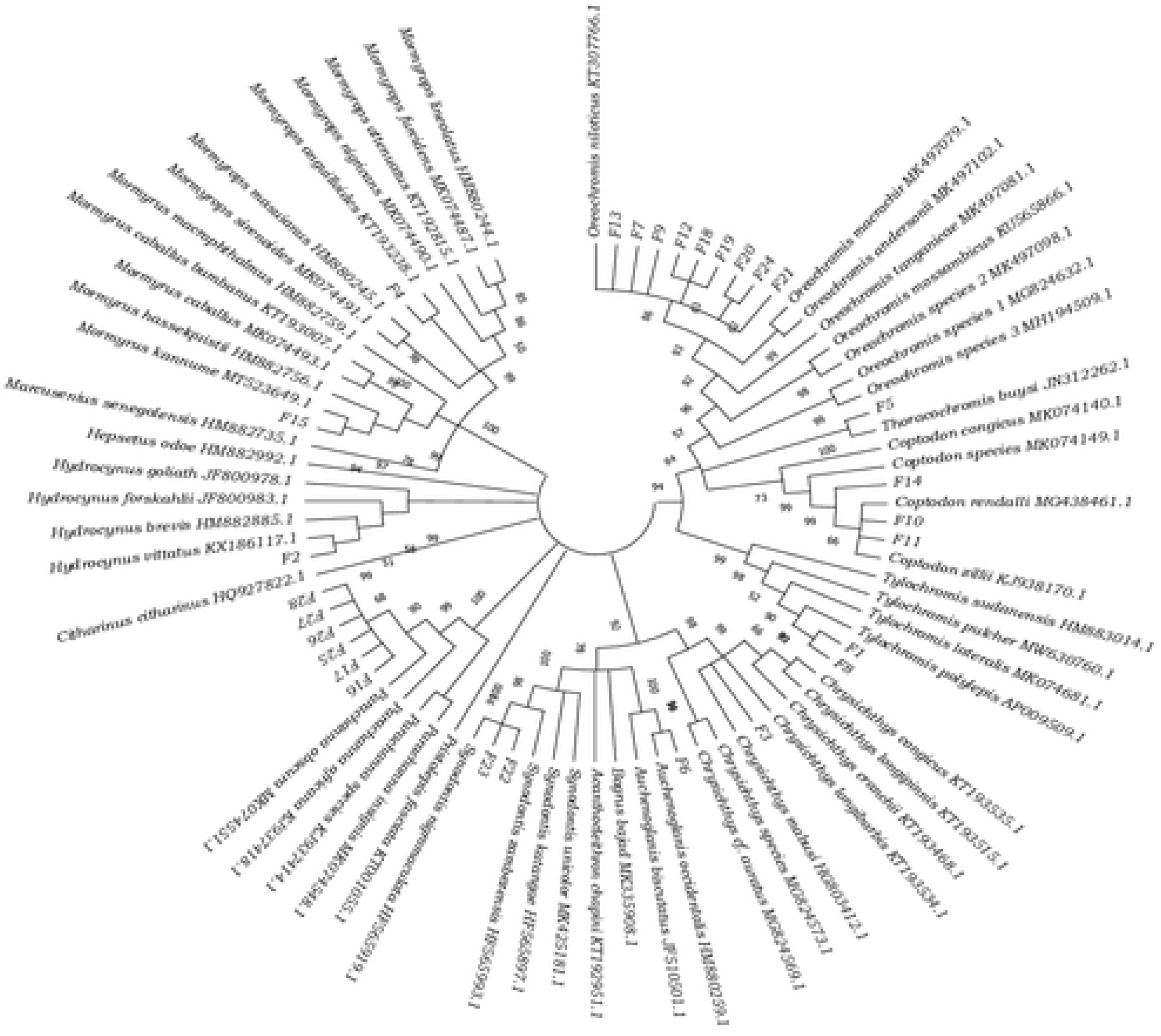
Consensus tree derived from COI sequences, using the Kimura 2-parameter model and 1,000 bootstrap replicates. All positions below 95% site coverage were removed, applying the incomplete deletion option. Analysis involved 84 nucleotide sequences, with 518 positions in the final dataset.

The *Oreochromis* sequence divergence values ranged from 0% to 10.5%. The highest pairwise comparison was between *O. tanganicae* and *Oreochromis* species 1 & 3 at 10.5%. All *Oreochromis* samples (F7, F9, F12, F13, F18, F19, F20, F21 & F24) provided the lowest divergence value of 0% for F7 and F13 and 0.2% for all other *Oreochromis* specimens against *O. niloticus*, showing a close relationship (S3 Table). The *Tylochromis* COI sequence divergence values ranged from 0.4% to 11.8% for the two representative samples/specimens. Both F1 and F8 specimens produced the lowest value of 0.4% relative to *T. polylepis.* This intraspecies COI sequence divergence value showed a close relationship to *T. polylepis*, a species that has never been captured by official DoF surveys and stock assessments. Specimen F5 showed a lowest sequence divergence value of 1% to *T. buysi* (Table S3). The *Coptodon* COI sequence divergences ranged between 0% and 12.4%. Specimens F10 and F11 provided the lowest divergence value of 0%, whereas specimen F14 provided 0.2 against *C. zillii*, showing a close relationship. Meanwhile, all the three specimens showed divergence values between 0.2% and 0.4% against *C. rendalli* (S3 Table).

### Channidae

All *Parachanna* specimens (F16, F17, F25, F26, F27 & F28) formed a well-defined group, with the best match to *P. obscura* MK074551.1 and similarity of 99.85% (Fig 2) The *Parachanna* COI sequence divergence values ranged between 0.2 – 11.2%, for all *Parachanna* specimens F16, F17, F25, F26, F27 and F28, with the lowest divergence value of 0.2% against *P. obscura*. Although *P. africana and P. insignis* showed a close relationship, they are clearly distinct (S3 Table).

### Alestidae

Specimen F2 with a lower similarity value of 96.69% appeared to be the most similar to *H. vittatus*, KX186117.1 (Fig 2). The *Hydrocynus* COI sequence divergence values ranged from 3.7% to 20.8%. Specimen F2 had the lowest divergence value of 3.7% compared with *H. vittatus* (S3 Table). The intraspecific divergence value of 3.7% is relatively high compared to the expected <2. This can be attributed to possible challenging taxonomies or misidentifications [57].

### Bagridae

Specimen F6 with a similarity identity of 99.69% was grouped with *A. occidentalis* HM880259.1 (Fig 2). The *Auchenoglanis* COI sequence divergence of 0.4% was the lowest divergence value for the specimen F6 against *A. occidentalis* (S3 Table).

### Claroteidae

Specimen F3 with a similarity value of 99.83% (Table 2) showed close relation to *C. mabusi* HG803412.1. Specimen F6 with a similarity value of 99.69% grouped with *A. occidentalis* HM880259.1 (Fig 2). The *Chrysichthys* COI sequence showed a divergence value of 0.2% when compared to *Chrysichthys mabusi* (S3 Table).

### Mochokidae

Specimens F22 and F23 with similarity values of 99.85% and 99.70%, respectively, were grouped with *S. nigromaculata* HF565919.1 (Table 2; Fig 2). The *Synodontis* COI sequence divergence ranged between 0.2% and 12%. The two specimens F22 and F23 displayed the lowest divergence value of 0.2% compared with *S. nigromaculata* (S3 Table).

### Mormyridae

Specimen F4 with a similarity identity of 99.54% was grouped with *M. anguilloides* KT193238.1, whereas sample F15 had a similarity of 98.46% with *M. kannume* MT523649.1 (Fig 2). This is the first record of *M. kannume* within the Mweru-Luapula fishery. The *Mormyrops* COI sequence divergence value of 0.4% was the lowest divergence value for the sample F4, when compared to *M. anguilloides*. The *Mormyrus* COI sequence divergence value of 2% was the lowest divergence value for specimen F15, when compared to *M. kannume* (S3 Table).

## Discussion

This study is the first to assess freshwater fish diversity of the Mweru-Luapula (ML) fishery, showing how DNA barcoding can complement survey data and morphological assessments, and tracking of invasive species as well as detection of introgression. The study further showed native species relationships based on phylogenetic trees. All *Parachanna* fin clips collected were identified as *P. obscura* [8], affirming this as the invasive species of the Mweru-Luapula fishery. A potentially new *Parachanna* species (*Pa*. sp. D. R. Congo, BIN AAF7843) observed by Conte-Grand et al. [25] could not be detected from the collected and analysed *Parachanna* specimens.

The study has demonstrated the strength and validity of COI barcodes as illustrated by Ward et al. [58] and Appleyard et al. [5] in identifying species using specific primer sets. Iyiola et al. [59] also tested the power of DNA barcoding by detecting cryptic diversity and analysis of intraspecific phylogeographic structure in the freshwater fishes of Nigeria. All 28 specimens studied were successfully amplified and sequenced. Sequences generated were used for performing BLAST searches on GenBank. This assemblage of *Oreochromis* species signified the strength of COI gene in suitably grouping species together, as illustrated by Tripathi [60]. All field morphologically identified *Oreochromis macrochir* were matched to a single species clade of *O. niloticus*. After identifying the mosty likely species, five species were a match with the phenotypical field identification and a further 22 samples were matched to the same genus level. Only one species was a complete mismatch. The barcode assignment or identification of the five species, *T. buysi, O. niloticus, T. polylepis, C. zilli* and *M. kannume* being reported for the first time in Mweru-Luapula fishery affirms the value of DNA barcoding. This was also observed by Sogbesan et al. [47], when the team first reported the presence of a sub-species, *Sarotherodon galilaeus boulengeri* in the Nigerian freshwater of Lake Geriyo. *Thoracochromis buysi*, a sister group to *Serranochromis* species, and *M. kannume* and *C. zilli* are native to Congo Basin [61] and could have moved into Mweru-Luapula fishery through migration, since the fishery is connected to the Congo Basin. *Oreochromis niloticus* could have escaped from fish culturing cages into the fishery from fish farmers keeping it illegally [35,36]. However, probable sources of *T. polylepis* into the fishery are yet to be established. These introductions were caused by the transfer of cichlids between freshwater lakes as noted by Dieleman et al. [62].

BLAST search results showed a success rate of the ‘best match’ at 100% for four specimens. These results were similar to studies done by Lakra et al. [63] and Shen et al. [64]. With the matching success range between 96.69 and 100%, the study showed efficiency in applying the DNA barcoding approach for identifying freshwater fish for the ML fishery, when compared to 90% matching success rate recorded in North America by April et al. [65], 93% in Canada by Hubert et al. [66] and 95.60% in North-Central Nigeria by Iyiola et al. [59]. This shows a reasonable identity based on BLAST, although it also demonstrates why BLAST comparisons need to be complemented by other methods to reliably identify species.

The phylogenetic analysis could not separate *O. andersonii* from *O. macrochir* into distinct groups, with a similar genetic divergence range of 1.4 to 3.8% as observed by Syaifudin et al. [67]. The consensus tree further supports what Bbole et al. [42] deduced in linking *O. andersonii* as a close relative to *O. macrochir.* This positioning shows the evolutionary relationship amongst the cichlids. Nevertheless, the presence of erroneous *Oreochromis* species sequences on GenBank database could have possibly influenced the placement of species on the phylogenetic tree as noted by Ordonez et al. [53] for suggesting EU752142, DQ856620, GU477624 and GU477627 to be inaccurate or a possible introgression. Results from this study are in line with Nwani et al. [68] for attributing detected ambiguities to taxonomic over-splitting, introgressive hybridization, recent radiation or incomplete lineage sorting. Harrison & Stiassny [69] also ascribed the loss of diversity in freshwater fish to unprecedented hybridizations from introductions of interrelated species. These observations stress the need for DNA barcoding to be tentatively embraced and form part of the routine monitoring tool, complementing officially recognised traditional methods in Zambia. However, for this recommendation to be actualised, it is essential to document sequence libraries for all fishes of Mweru-Luapula and other fisheries in the country.

The clustering of the collected COI sequences alongside the retrieved GenBank reference sequences for *O. niloticus*, *O. macrochir* and *O. andersonii* corroborates with findings by Falade et al. [70] in South-Western Nigeria and Iyiola et al. [59] in North-Central Nigeria. This clustering of our studied freshwater species also affirms the assertions by Blackwell et al. [71] that gene pools can readily be impacted by hybridisation between alien and native congeneric species. The tendency by *Oreochromis* species to hybridise, especially with tilapia was elaborated by Angienda et al. [72] and Deines et al. [73] as the reason why cichlids cannot easily be distinguished based on morphological identifications. There is a high probability of the presence of hybrids in the studied populations of the Mweru-Luapula fishery. This assertion supports the tendency by invasive *O. niloticus* to introduce genes into numerous congeneric populations with weak equals as observed by Fatsi et al. [74].

The two distinct lineages identified in the *Coptodon* phylogenetic analyses show a possible likelihood of cryptic lineage diversity within the freshwater fishes. This is similar to what Iyiola et al. [59] established when analysing the freshwater *Synodontis intermedius*. The placement of *C. zillii* KJ938170.1 and *C. rendalli* MG438461.1 clustered with the collected COI sequences supports the phylogenetic placement of these species as close relatives as observed by Bbole et al. [42]. The distinction of closely related species demonstrates the efficacy of the COI gene, as was propagated by Ude et al. [75]. However, this ambiguity in *Coptodon* species assignment observed demonstrates the need for a high-quality sequence reference and appropriately assigned voucher specimens, which are currently difficult to obtain from both GenBank and BOLD databases as observed by Diaz et al. [9] and Iyiola et al. [59].

Identification mismatches were encountered between species assigned using field morphological methods and the DNA barcoding approach. The mismatches may represent phenotypic variation and thus field misidentification [53]. On the other hand, the mismatches observed can potentially be attributed to hybridization and introgression that would have led to morphological features being similar to one species but the DNA sequence of a different taxon [76,77]. This explains mismatches or misidentifications encountered between field identifications and DNA barcodes assignment. The study demonstrates the challenge taxonomists face when they solely depend on morphological characteristics to identify species. Challenges with morphological field identification emphasizes the importance of using combined molecular and morphological approaches, as highlighted by Iyiola et al. [59]. These challenges are exacerbated by a lack of key guides for most Southern African freshwater fishes. The possible mismatches of *T. polylepis* for *T. mylodon, T. buysi* for *A. macrophthalmus, O. niloticus* for *O. macrochir; C. zillii* for *C. rendalli, M. kannume* for *M. longirostris* and *S. nigromaculata for S. katangae* from field identification validates the observations made by Erwin & George [7] stressing the limitations of solely using visual identifications due to convergent morphological characteristics in some species. In line with Becker et al. [78], a precautionary approach should be incorporated in addressing taxonomic mismatches arising from low quality sequences submitted to the GenBank database.

The low divergence values observed from the analysis of the collected samples against *O. niloticus* show minimal genetic differences within species, while the high genetic distance of 10.40% observed, when compared to *O. tanganicae* supports the argument that the species are distinct. This is similar to what Ward et al. [58] suggested that due to possible species fusion, distinct species may have identical COI sequences in some cases.

## Conclusion

Our study has provided 28 COI barcode sequences, including 15 new species or new records of fish species of the ML fishery. The identification of five collected samples to species level and 22 to genus level underscores the reliability of DNA barcodes. The analysis of randomly collected fin clips has also detected *P. obscura* [8] as the snakehead fish that has invaded the ML fishery, and not the new *Parachanna* species observed in the Congo Basin [25]. We therefore recommend further detailed sampling, collection and analysis of samples on a large scale. By submission of our findings in this study, we have contributed to the building of a DNA reference barcode database of Zambian native fish fauna, thereby increasing the resources on GenBank and BOLD.

## Acknowledgements

Gratitude is extended to the Zambia Aquaculture and Enterprise Development Project (ZAEDP) under the Department of Fisheries in the Ministry of Fisheries and Livestock (MFL) for the provision of field staff, transport, gears, boats and other equipment and materials for use during field data collection. We are also highly indebted to Dr. Stephen Koblmüller (University of Graz and ABOL, Austria) and Dr. Herman Chambaro for their valuable insights and contribution towards data analysis. Last, but not the least Mr. Obed Chanda, under the Ministry of Agriculture is recognized for the production of maps showing the location of the sampled sites.

## Supporting information

**S1 Table Gill net survey fleets**

**S2 Table Reference species sequences for ML fishery obtained from NCBI GenBank database**

**S3 Table Intraspecies and Interspecies pairwise genetic distances (%) of COI sequence data of the Mweru-Luapula fishery using Kimura-2-parameter.**

**S1 Fig The original phylogenetic tree showing comparative positions of the 28 specimens and retrieved GenBank sequences based on Maximum Likelihood consensus tree (inferred from 1,000 replicates of COI gene), before collapsing the tree to cater for >50% bootstrap values.**

